# *In Silico* Engineering of Stable siRNA Lipid Nanoparticles: Exploring the Impact of Ionizable Lipid Concentrations for Enhanced Formulation Stability

**DOI:** 10.1101/2024.09.25.614899

**Authors:** Malay Ranjan Biswal, Sudip Roy, Jayant K Singh

## Abstract

Lipid nanoparticles (LNPs) are crucial in advancing the delivery of RNA-based therapeutics within the domain of gene therapy. A comprehensive understanding of their formation and stability is critical for optimizing the clinical efficacy of LNPs. This study systematically investigates the influence of concentration variations of positive and neutral ionizable lipids - specifically, 2-[2,2-bis[(9Z,12Z)-octadeca-9,12-dienyl]-1,3-dioxolan-4-yl]-N,N-dimethylethanamine (DLinKC2-DMA) and 1,2-Dioleoyl-sn-glycero-3-phosphoethanolamine (DOPE) - along with cholesterol and polyethylene glycol, on the formation of LNPs and encapsulation of small interfering RNA (siRNA). Utilizing coarse-grained classical molecular dynamics (MD) simulations with a system size matching experimental range (approximately 0.6 million beads), we conduct a comparative analysis and offer mechanistic insights into siRNA formulation within LNPs containing positive and neutral DLinKC2-DMA. We found that the LNPs with positive ionizable lipids encapsulate more than twice the siRNA compared to the LNPs with neutral ionizable lipids. In addition to the formation of LNPs, our study extends to the forces governing siRNA escape from LNPs, employing steered molecular dynamics simulations. The force experienced by siRNA to cross the LNP lipid layer containing positive ionizable lipids was 400kJ/mol/nm more than that of neutral ionizable lipids, suggesting the encapsulation is more favorable with positive ionisable lipids.

## Introduction

RNA interference (RNAi) or post-translational gene silencing plays important practical applications in gene therapy. Certain nucleic modalities like siRNA and mRNA are inserted into the tissue to regulate gene expression during this process.^1^ After the first demonstration of the potential of siRNA therapeutics, where intravenous injection of Fas siRNAs protected mice from liver fibrosis, drug development has been rapid.^2^ Due to their ability to downregulate gene expression guided by sequence complementarity, siRNA presents an extraordinary potential to block the synthesis of disease-causing proteins. siRNA helps against diseases caused by gene overexpression, mRNA on the other hand tackles diseases with insufficient protein production, which was observed in mRNA vaccination during the COVID-19 pandemic.

RNA-based drugs have the potential to be the future of drugs and medicine as there are many advantages over traditional medicines. It is easy to synthesize, customize, and modify mRNA and siRNA. In COVID-19, the mRNA vaccine was the first to reach the phase I trial in a few months.^3,4^ These vaccines are highly targeted as they express specific antigens and induce a direct immune response.^5,6^ Despite multiple appealing features, there are many challenges in vivo delivery, primarily because of the (1) instability of naked and unmodified RNA due to enzymatic degradation by ribonuclease present in the blood,^7^ (2) the highly negative charge with large size of mRNA creates insufficient intracellular delivery,^8,9^ (3) poor stability and pharmacokinetics.^10^

Two major technologies developed to overcome these challenges - (a) GalNAc^11^ (b) liposomes or lipid nanoparticles (LNPs).^12–14^ GalNAc is an amino acid derivative of galactose, forms conjugate with siRNA and binds to the Asialoglycoprotein receptor which is expressed predominantly in hepatocytes.^15^ On the other hand, vesicles like liposomes and LNPs carry not only RNAs but several other hydrophobic drugs^16^ and cellular materials, like proteins,^17^ lipids^18^ and DNA,^19^ from source to distant sites via biofluids. Studies have shown the vesicle formation using various phospholipds, amphiphiles, surfactants, etc.^20–22^ The vesicles protect nucleic acids and other biological molecules from degradation in *in vivo*, and deliver them to the targeting sites without interfering with immune activation. These liposomes and nanoparticles have been used in the treatment of cancer^23^ and vaccines.^24,25^ Many of these liposomes have been used as a carrier in clinical trials to deliver anticancer, anti-inflammatory, antibiotic, antifungal, anesthetic, and other drugs and gene therapies, as they help the biomolecules to cross the negative potential cell membrane barrier. These LNPs effectively deliver nucleic acids such as RNA or DNA-based drugs by encapsulating them inside the micelles or vesicles. In 2018, first siRNA drug, patisiran was approved, and later givosiran, lumasiran, etc. were also used as siRNA drugs.^26^ Also, two authorized COVID-19 vaccines, mRNA-1273^27,28^ and BNT162b2^29^ use LNPs to deliver antigen mRNA.

A typical LNP for RNA-based drug delivery contains mainly four types of lipids - ionizable lipids, polyethylene glycol (PEG)-functionalized lipids (PEG-lipids), cholesterol and other phospholipids such as phosphatidylcholine or phosphatidylethanolamine.^30–32^ Ionizable lipids are similar to that natural lipids except for the presence of ionizable head groups that become positively charged in an acidic environment and stays neutral in the basic environment instead of zwitterionic or anionic head group of natural lipids; because of which they are less toxic in the bloodstream compared to non-ionizable cationic lipids.^33^ Due to the presence of these positively charged lipids, the nucleic acids get stabilized, and resistance to nucleic degradation is increased. Apart from the ionizable lipids, other lipids molecules such as PEG-lipids, phospholipids and cholesterols improve stability, delivery efficacy, tolerability and biodistribution. PEG-lipids can affect the size of LNP, reduce particle aggregation, prolong the blood circulation time of nanoparticles and conjugate specific ligands to the particle for targeted delivery.^31,32,34–36^ Phospholipids help in the stabilization of LNPs and also facilitate endosomal escape.^37,38^ The presence of cholesterol also stabilizes LNPs by modulating the membrane integrity and rigidity. It also affects the delivery efficacy and biodistribution of nanoparticles.

LNPs are typically produced through the microfluidic mixing method, involving the passage of RNA and lipids dissolved in an acidic organic solvent through microfluidic channels.^39^ Within this environment, the negatively charged RNA interacts with ionizable lipids, which transition to a positively charged state in the low pH surroundings, facilitating the formation of small lipid droplets that coalesce into LNPs. Following production, the LNPs are typically stored under neutral or physiological conditions.

Many studies have addressed the encapsulation of RNA-based drugs in LNPs, and researchers have narrowed down various lipids that are essential to formulate RNA,^40^ but the exact mechanism of how these RNAs are encapsulated in the LNPs system still needs to be understood. Several LNPs developed in the last few years are either in the preclinical phase or could never reach human trials. Thus, understanding the composition and stability of various LNPs is necessary to deliver siRNA vaccines while simultaneously looking for delivery safety. An earlier study utilized coarse-grained molecular dynamics to investigate the encapsulation of small DNAs within lipid nanoparticles.^41^ Recently, Trollmanm and Böckmann^42^ elucidated the lipid arrangement influenced by the protonated and deprotonated states of ionizable lipids within a lipid bilayer. Their primary focus was on understanding lipid arrangements with different protonation states, both in the presence and absence of RNA, which they observed in bilayers. Their findings in full scale LNP indicated that mRNA-free LNPs with deprotonated ionizable lipids lacked water pockets. Inspired by their work, we advanced further by exploring the variation in the formation and encapsulation of siRNA using various concentrations of ionizable and phospholipids. However, Paloncýová *et. al.*^43^ demonstrated differences in mRNA encapsulation within LNPs containing neutral and positively charged ionizable lipids. Due to the limited number of lipids, siRNAs are not fully encapsulated. Consequently, when the ionizable lipids are deprotonated, siRNAs are expelled from the LNPs. This raises concerns about whether RNAs remain intact within LNPs in the bloodstream, which has a high pH condition. Apart from LNP formulation, coarse-grained molecular dynamics were also employed to investigate DNA endosomal escape in cationic liposome complexes .^44^

In this study, we studied the LNP formulation with varying concentration and charges of ionizable lipids from molecular simulation. The work aims to compare the encapsulation of siRNA in LNP containing positive ionizable lipids and LNP containing neutral ionizable lipids during the LNP formation process using coarse-grained molecular dynamics (MD) simulation. These types of LNPs resemble the encapsulation of siRNA in acidic and neutral mediums, respectively. We varied the concentration of ionizable lipids (DLinKC2-DMA) in the LNPs into three categories: low concentration, medium concentration, and high concentration. To the best of our knowledge, this study is one of the first attempts to showcase how computational modeling and simulation can help design RNA-based drug delivery systems for future pharmaceutical industries.

## Modelling and computational details

We aim to study the siRNA formulation in lipid nanoparticles within the experimental size range. However, using an all-atom structure can be computationally challenging and costly when simulating large lipid vesicle systems, such as those in the 10-150 nm range.^45^ To address this, we have employed a coarse-graining approach. Coarse-graining helps capture the physics at a large length scale by reducing degrees of freedom, making it feasible to simulate large-scale bio-vesicles. Given the complexity of lipid vesicle systems and the requirement for lengthy simulation times, coarse-grained MD modeling has proven advantageous.^46^

In addition to investigating the self-assembly of lipid nanoparticles, we also wanted to understand the force experienced by siRNA to cross the membrane layer of the vesicle using steered molecular dynamics (SMD) simulation. This will help us to comprehend the escape of siRNA from LNPs and the stability of LNPs. The study is limited to a layer of membrane and its interaction with siRNA, and therefore, it is not necessary to simulate whole of the LNP. Instead, a bilayer and one siRNA molecule can be simulated, reducing the system size by ten times. Although, performing SMD in coarse-grained system will be faster, it will lead to extra complexity like low degrees of freedom and elimination of fine interaction details. Since the system size was reduced, we performed SMD with an all-atom structure to get better insight, high degrees of freedom and detailed interaction.

### Coarse-graining and system preparation

Martini coarse-grained (CG) version 2 was employed for molecular dynamics simulation. For CG mapping, four heavy atoms were represented as one single CG bead (“four-to-one”) as per the Martini philosophy. For smaller beads three-to-one and two-to-one are also implemented, if required. For the CG Martini mapping of the positively charged lipid molecule DLinKC2-DMA (DLKC2), an all-atom structure was manually coarse-grained, following the CG-Martini guidelines,^47^ and the methods followed by Leung *et, al* (Figure 1).^41^ The same processes were repeated, replacing the protonated DLKC2 (with the head group containing charged Qd type) with neutral DLCK2 (with the head group containing Na type). An all-atom model of siRNA was obtained from the PDB id: 2F8S. The all-atom RNA model was then coarse-grained using the “martinize-nucleotide python script”.^48^ For DOPE and cholesterol (CHOL), the CG structures were obtained from Marrink *et, al*.^49^ Finally, the PEGylated lipid molecule was prepared using the DOPE as lipid moiety and available CG PEG polymers with sixteen ethylene glycol CG Martini beads.^50^ The CG maping of the structures are given in Supplementary Figure S1-S4. To neutralise the system, sodium (NA) or choride ions (CL) were used based on the net charge of the lipids and siRNA. The charges for each protonated DLKC2 and siRNA are +1 and −42 respectively.

**Figure 1:**
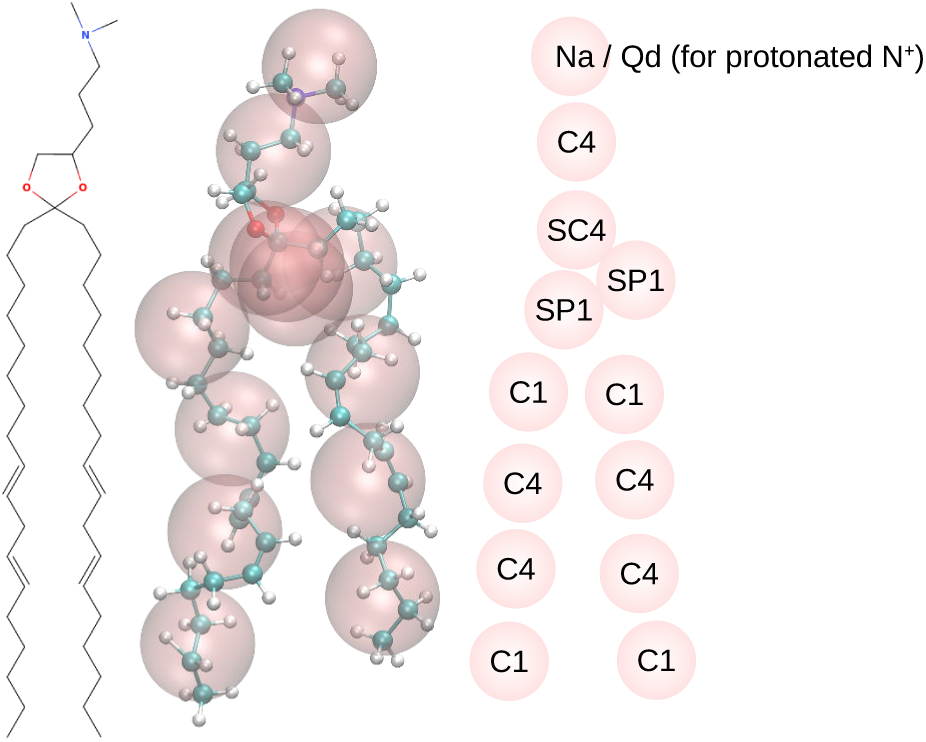
CG Martini mapping of DLinKC2-DMA. The bead types are according to Martini version 2 convention. For neutral lipid, the head group type is Na, and Qd for positive lipid.

As represented in Figure 2, at first, three types of small boxes of size 8×8×8nm^3^ were created with varying DLKC2 and DOPE concentrations (mentioned in Table 1). In each of these boxes, two siRNAs, 576 cholesterol were also placed. In our system, we varied only DLKC2 and DOPE keeping the total weight% of all the lipids same. The low, medium and high concentration referred in the Table 1 is with respect to DLKC2. The N/P ratio corresponding to low, medium and high concentration are 1.6, 6.3, and 7.3, respectively. Studies have reported that varying the N/P ratio affects the encapsulation efficiency of mRNA and siRNA.^51^ After adding 4030 water beads, including 10 percent non-freezing water beads and the appropriate amount of Na or Cl ions to neutralize the systems, the systems were self-assembled, starting from random configurations, using GROMACS 2019^52^ as mentioned in the computational details section. As the main important electrostatic interaction considered here is between siRNA and lipid molecules, we considered normal water beads instead of polarizable water beads. Use of polarizable water beads will increase the system size, and add complexity. After the self-assembly of each of the small building blocks for 1 *µ*s, larger systems were created by spatially translating the block. These larger systems were 8 times the smaller building blocks, repeated 2 times in all three directions. The larger system then kept in the center of a bigger box of size 40×40×40 nm^3^. Then 1152 PEG-lipids molecules and additional 375000 water beads (including 10% non-freezing water beads) were added. The composition of the larger system is given in the Table 1. A further 5 *µ*s of molecular dynamics simulation was performed to get a spherical shaped LNP. Each case was repeated twice to get an average estimate for each case, i.e. total 12 systems were simulated.

**Figure 2:**
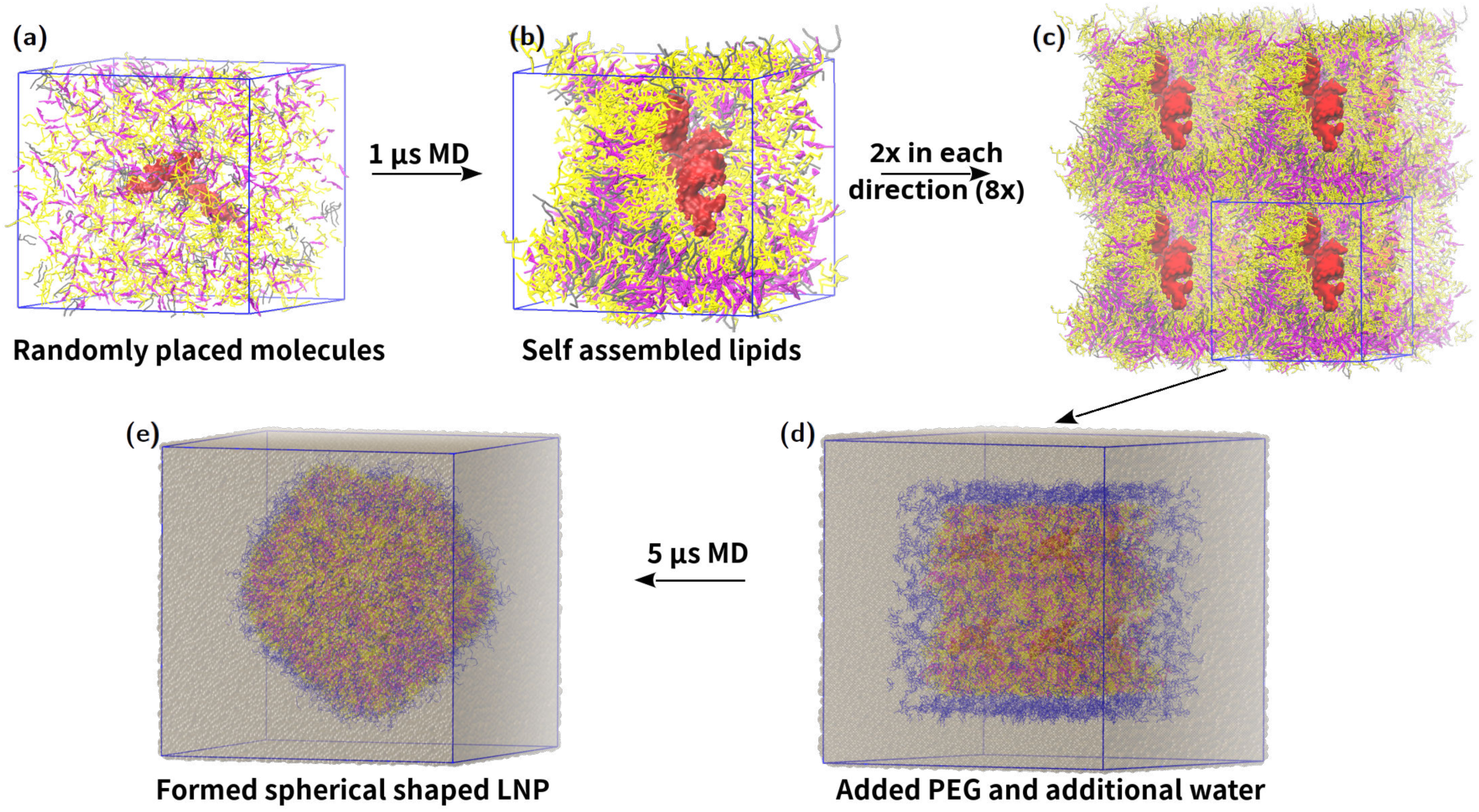
Steps followed for coarse grained MD simulation. (a)All the molecules are placed randomly in a box, (b)Self assembled structure of lipids is seen after 1 *µ*s of MD simulation, (c)box is repeated 8 times, i.e, 2 times in each direction, (d)the repeated structure was placed in a bigger box, and PEG-lipids, additional water and ions were added, (e)After 5*µ*s of MD simulation spherical shaped LNP was obtained. DLKC2 is shown in yellow, DOPE in gray, cholesterol in pink, PEG-lipids in blue and RNA in red; water is not shown for clarity in (a)-(c). In (d) and (e) water is grey.

**Table 1:** Three different types of concentration were taken with respect to DLKC2, in the small building block. The medium concentration was based on an earlier work.^41^ The weight percentage is given in the bracket adjacent to the number of the lipids.

|  | DLKC2<br>(positive/<br>neutral) | DOPE | CHOL | Lipid ratio | RNA | Water |
| --- | --- | --- | --- | --- | --- | --- |
| Low Concentration | 144 (8.6) | 576 (40.0) | 576 (21.5) | 1:4:4 | 2 (2.6) | 4030 (27.0) |
| Medium Concentration | 576 (35.5) | 144 (10.3) | 576 (22.1) | 4:1:4 | 2 (2.6) | 4030 (27.8) |
| High Concentration | 648 (40.1) | 72 (5.2) | 576 (22.2) | 4.5:0.5:4 | 2 (2.7) | 4030 (27.9) |

**Table 2:**
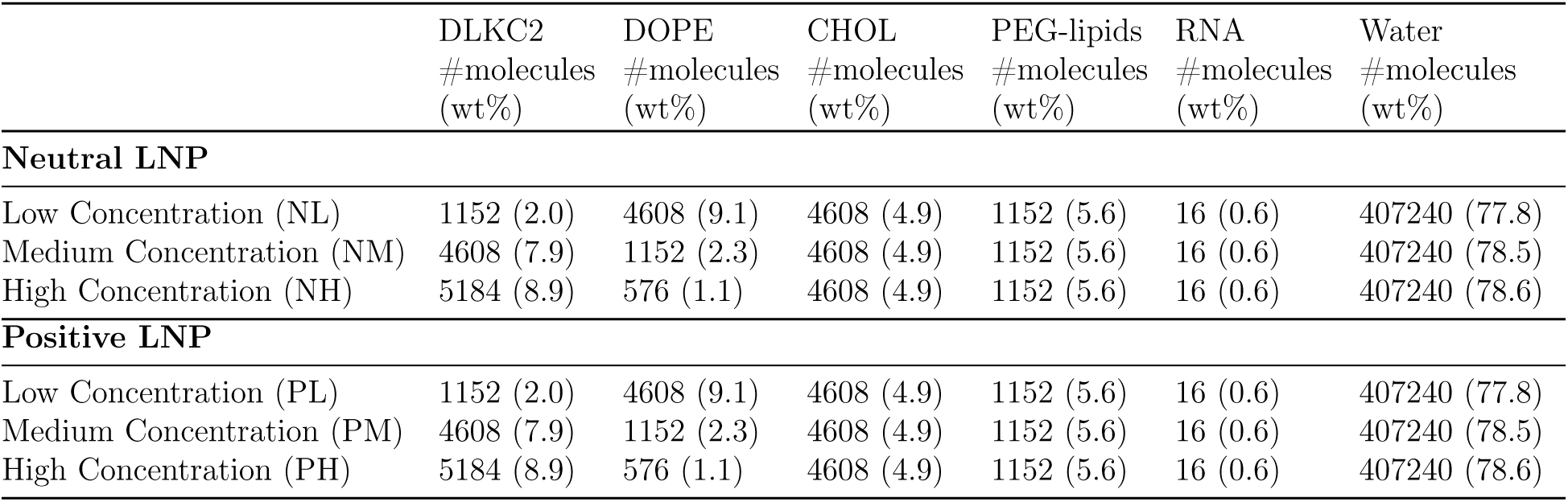
LNP compositions after creating the larger system by repeating the box two times in all direction. The weight percentage is given in the paranthesis.

**Table 3:**
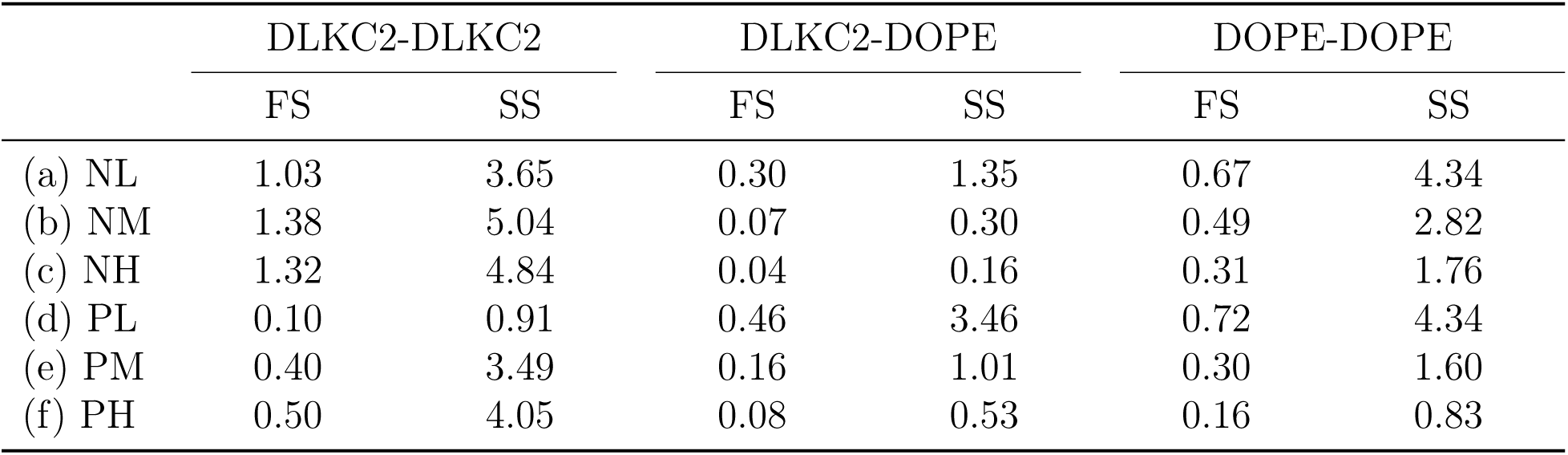
Coordination number of first shell (FS) and second shell (SS) of various systems from Figure 6.

### Computational details

All the molecular dynamics simulations were performed using GROMACS version 2019.^52^ The system was minimized with the steepest descent minimization. For neighbor searching, a verlet cut-off scheme was used with frequency to update the neighbor list set to 10. Reaction field electrostatics with a Coulomb cut-off of 1.2 nm and van der Waal’s cut-off of 1.2 nm were used. A value of 15 was used for the relative dielectric constant. A temperature coupling was set to velocity rescale with reference temperature 310 K and time constant for coupling 0.1. For NPT simulation, isotropic Parrinello-Rahman pressure coupling^53^ of 1 bar was used with a time constant of 10 ps. A periodic boundary condition in all directions was used.

### Models and computational details for steered molecular dynamics study

To understand the force experienced by siRNA to cross the lipid layer of LNPs, steered MD study was performed. An all-atom bilayer system was prepared with a total of 144 cationic DLKC2, 36 DOPE, and 144 cholesterol as lipids in each of the upper and lower layers of the bilayer. The above-mentioned siRNA was placed 4 nm above the upper layer, and then the system was solvated and neutralized with the required amount of water and ions. The starting structure for the SMD is given in Figure S5. After system preparation, NVT and NPT equilibrations were performed for one nanosecond. After the equilibration, steered MD calculation was performed using GROMACS 2019. ^52^ A harmonic potential of force constant 600 kJ/mol/nm^2^ was applied between the center of mass of siRNA and a virtual particle moving along the negative z-direction with a constant velocity, 0.5 nm per ns. During this time, the center of mass of siRNA was pulled for 30 ns in the negative z-direction allowing to cross the lipid bilayer smoothly. The simulation was performed in NPT ensemble in a box size of approximately 13×13×33 nm^3^. A semi-isotropic pressure coupling with Parrinello-Rahman barostat^53^ was used. The process was repeated with neutral DLKC2 lipids keeping all other configuration same.

## Results and discussion

### Formation of LNP with coarse-grained simulation

Figure 3 presents the snapshot of the self-assembled lipids after 1 *µ*s of coarse-grained molecular dynamics simulation for different concentrations of DLKC2. In Figure 3(a), 3(b) and 3(c), depicting low, medium and high concentration of DLKC2, respectively, the lipid layer and water are seen separated and siRNAs are wrapped around a lipid bilayer. These self-assembled structures were used for creating larger systems for a more extended MD simulation, as mentioned in the methods section. Figure 4 show that after 5 *µ*s of MD simulations, a clear distinction of compartments was formed where siRNAs were encapsulated, confirming the formation of lipid nanoparticles. At low and medium concentration of neutral and positive DLKC2 (Figure 4(a), (b), (d) and (e)), the compartments exhibit a noticeable lipid bilayer, in contrast to high concentrations of DLKC2 (Figure 4(c), and (f)). In these instances, the LNPs failed to form a well-defined lipid bilayer between these pockets and the outer surface. Earlier studies by Kulkarni et al., have shown that phospholipids are important in the formation of a stable LNP.^54,55^ Lower concentration of phospholipids creates low encapsulation efficiency.

**Figure 3:**
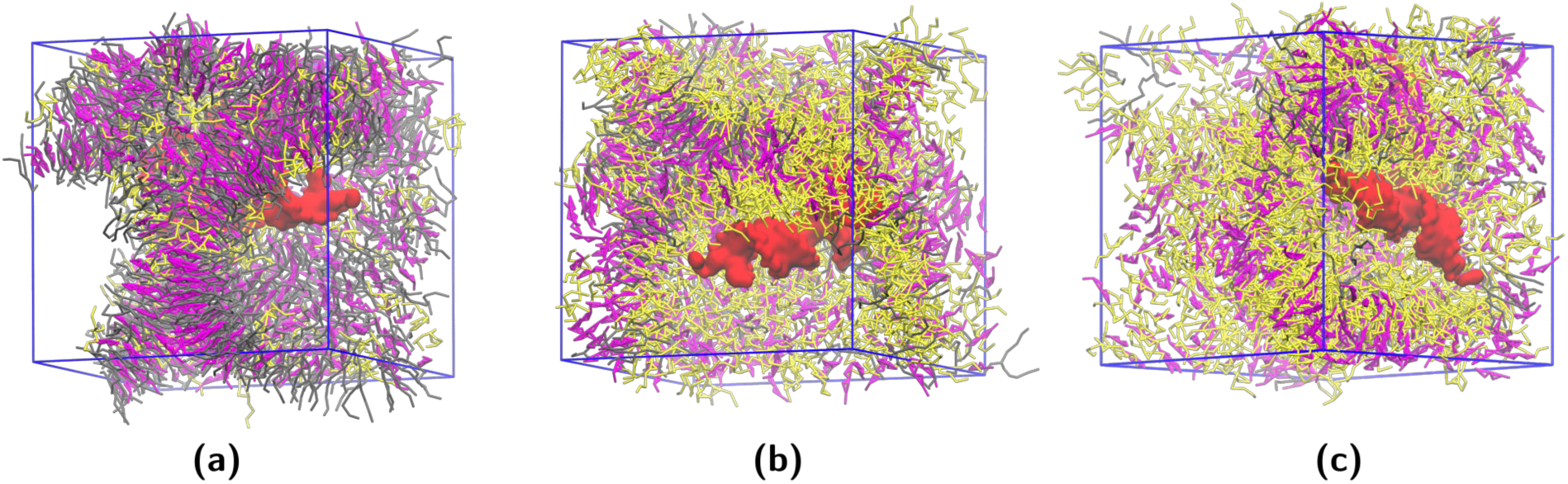
After 1 *µ*s of MD simulation shows a self assembly of lipids in the small building block for LNP. Three types of concentration of DLKC2 are shown ((a) low, (b) medium and (c) high, respectively). DLKC2 is shown in yellow, cholesterol in pink, DOPE in gray, lipid head group in cyan, and RNA in red; water is not shown for clarity.

**Figure 4:**
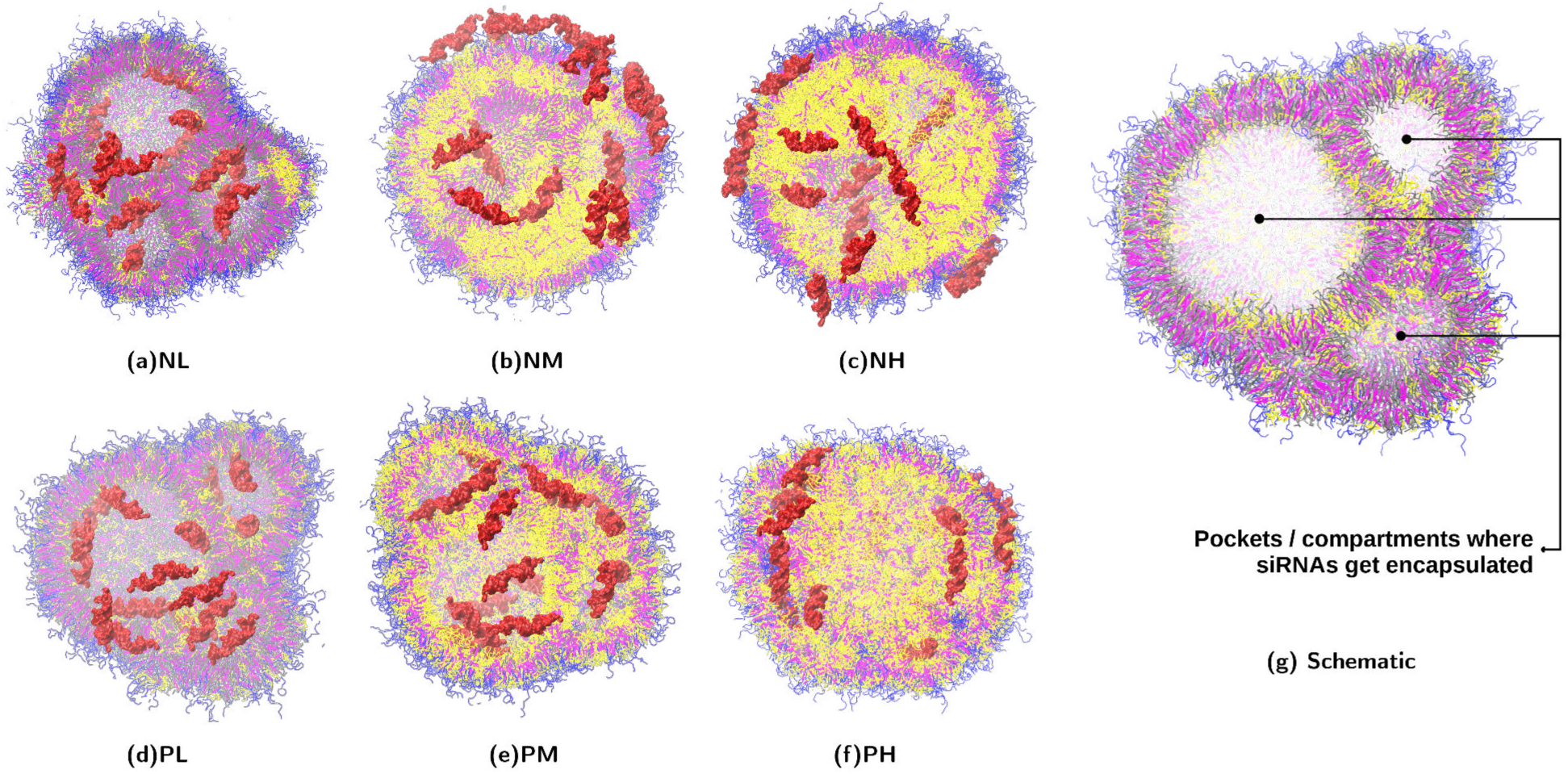
Self-assembled system after 5 *µ*s of simulation. (a)NL, (b)NM and (c)NH denotes Neutral DLKC2 in Low, Medium and High concentration; (d)PL, (e)PM and (f)PH denotes Positive DLKC2 in Low, Medium and High concentration. DLKC2 is shown in yellow, cholesterol in pink, DOPE in gray, and RNA in red; water is not shown for clarity. (g) A schematic of LNP is shown to represent compartments where siRNAs are encapsulated.

During the first 1 *µ*s, the PEG-lipids molecules got adsorbed onto the surface of the nanoparticles, where the hydrophilic parts were in the water. During simulation, the PEG-lipids molecules homogeneously got distributed on the outer layer of the LNP, and the LNP slowly transformed into a more rounded structure, with all the PEG-lipids molecules surrounding the outer layer. The diameter of the LNPs was roughly 35 nm, which is within the range of experimental observation.^56^ Inside the LNPs, the siRNAs were seen situated at various pocket-like structures. Each pocket was separated by a bilayer, where the encapsulated siRNA was attached to the lipid head groups. After repeating the simulation, we noticed similar behavior in both simulations. RMSD plot for the lipids in Figure S6 shows that lipid vesicle formation is stabilized after 4*µ*s in all the systems.

The number of siRNAs inside and outside the LNP was calculated and plotted in Figure 5. It was observed that, all the siRNAs are encapsulated in lipid nanoparticles with low and medium concentrations of DLKC2. Twelve (>0.15 weight%) out of sixteen siRNAs were seen encapsulated in case of high concentration of positive DLKC2. In contrast, the number was less than five in the case of neutral DLKC2, except with low concentration. In the nanoparticles with low concentrations of neutral DLKC2, all the siRNAs were captured inside the LNP with better compartments, similar to the cases with positive DLKC2. In all the systems, whether siRNAs were outside the LNP, the siRNA remained attached to the headgroup of the lipids in the bilayer.

**Figure 5:**
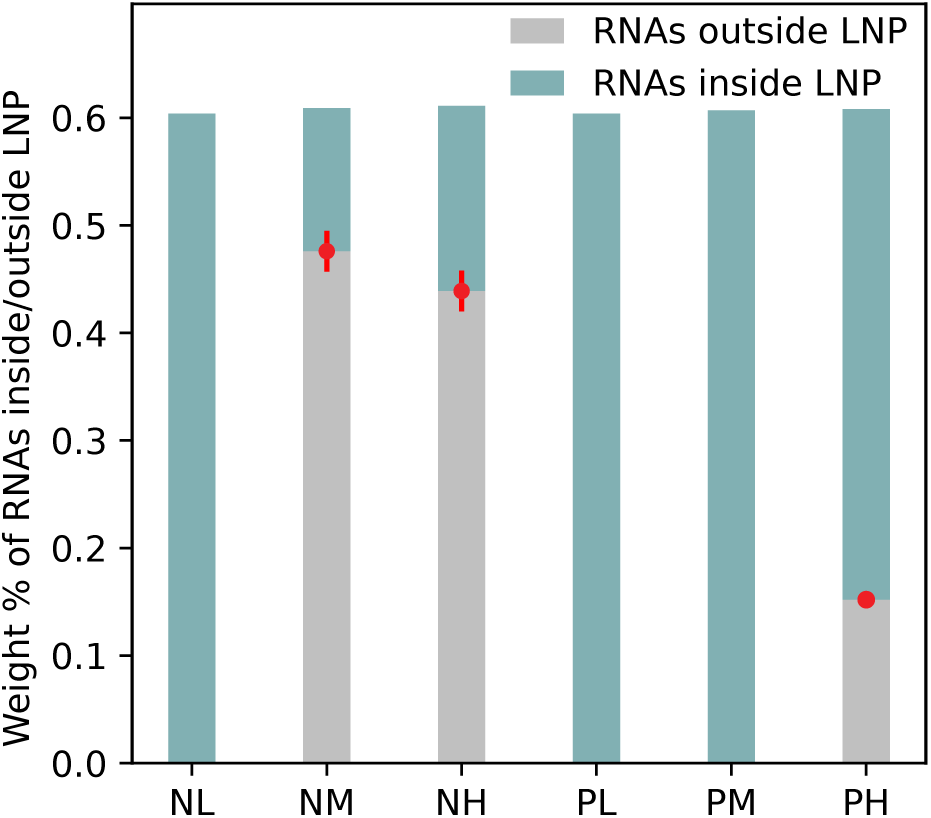
Weight percentage of siRNA captured inside the LNP in the systems after 5 *µ*s of MD simulation. The values are averaged over two repeated simulations. As metioned in the Table 1,NL, NM, and NH, represent neutral DLKC2 with low, medium and high concentrations in LNP. PL, PM, and PH, represent cationic DLKC2 with low, medium and high concentrations in LNP.

It was also observed that the siRNA molecules were better captured in systems with a higher number of compartments, in addition to the charges of ionizable lipids. The number of compartments formed varied depending on the concentration of neutral and positive DLKC2 in the LNPs. For instance, LNPs with low, medium, and high concentrations of neutral DLKC2 had four, two, and two compartments, respectively. Similarly, LNPs with low, medium, and high concentrations of positive DLKC2 had five, six, and two compartments, respectively. In the case of high neutral DLKC2 (NH) and high positive DLKC2 (PH) concentrations, both NH and PH had only two compartments, with one compartment containing two siRNAs and another containing three siRNAs. In the PH case, four siRNAs remained outside the LNP, while the rest seven siRNAs were encapsulated within two bilayers of the LNP. In any systems, a maximum of 5 siRNAs were seen encapsulated in a single compartment.

To understand the distribution of ionizable lipids and phospholipids in the nanoparticles, the radial distribution function of lipid head groups was calculated from last 1*µ*s trajectory as shown in Figure 6. A higher value of the g(r) at a closer radius was observed for a low concentration of DOPE. However, in Figure 6(a), a very high g(r) is seen with DLKC2-DLKC2, suggesting the ionizable lipids are closer to each other compared to other lipids. This is further illustrated in Figure 7. Compared to positive LNPs (Figure 6(d), (e), and (f)), the DLKC2 in neutral LNPs (Figure 6(a), (b), and (c)) are more aggregated. As the ionizable lipid which are the major component in encapsulating nucleotides are locally aggregated (refer Figure 7), there are large areas in the LNPs where there are no ionizable lipids. Due to this there is a possibility that the encapsulated siRNAs can escape the LNP in the later stage, which affects the stability of LNP. Compared to the case of low concentration of postive DLKC2, aggregation is not seen (Figure S7). Similar aggregation tendencies of neutral DLKC2 was also observed in earlier molecular dynamics studies of bilayer membrane with DLKC2 and POPC lipid,,^57^ as well as studies on the arrangement of DLin-MC3-DMA ionizable lipids within DOPE and DOPC phospholipids.^58,59^ These findings also suggest that neutral ionizable lipids tend to aggregate, whereas positive ionizable lipids tend to integrate with phospholipids. Additionally, atomistic membrane bilayer simulations have revealed that higher concentrations of neutral ionizable lipids further promote aggregation.^60^ From Figure 8, the short-range LJ potential and Coulomb energy between RNA and lipids show more negative values in LNP with positive DLKC2 compared to LNP with neutral DLKC2. It is due to a stronger electrostatic interaction between negative RNA and positive DLKC2. The higher the concentration of positive DLKC2, the stronger the interaction between RNA and lipids. Also, comparing between LNPs with neutral DLKC2, stronger interaction is seen with a higher concentration of DOPE (Figure S8). The presence of the high concentration of DOPE phospholipids, could have lead to the encapsulation of siRNA. The Martini model of DOPE contains both positive (quaternary ammonium) and negative (phosphate) beads. This is also seen in Figure 4, where the RNA molecules are bound to the hydrophilic headgroup of the lipid molecules. Similar observation was found in the earlier studies that the lipid polar headgroup interact with the negatively charged nucleotides.^41,43,54,55^

**Figure 6:**
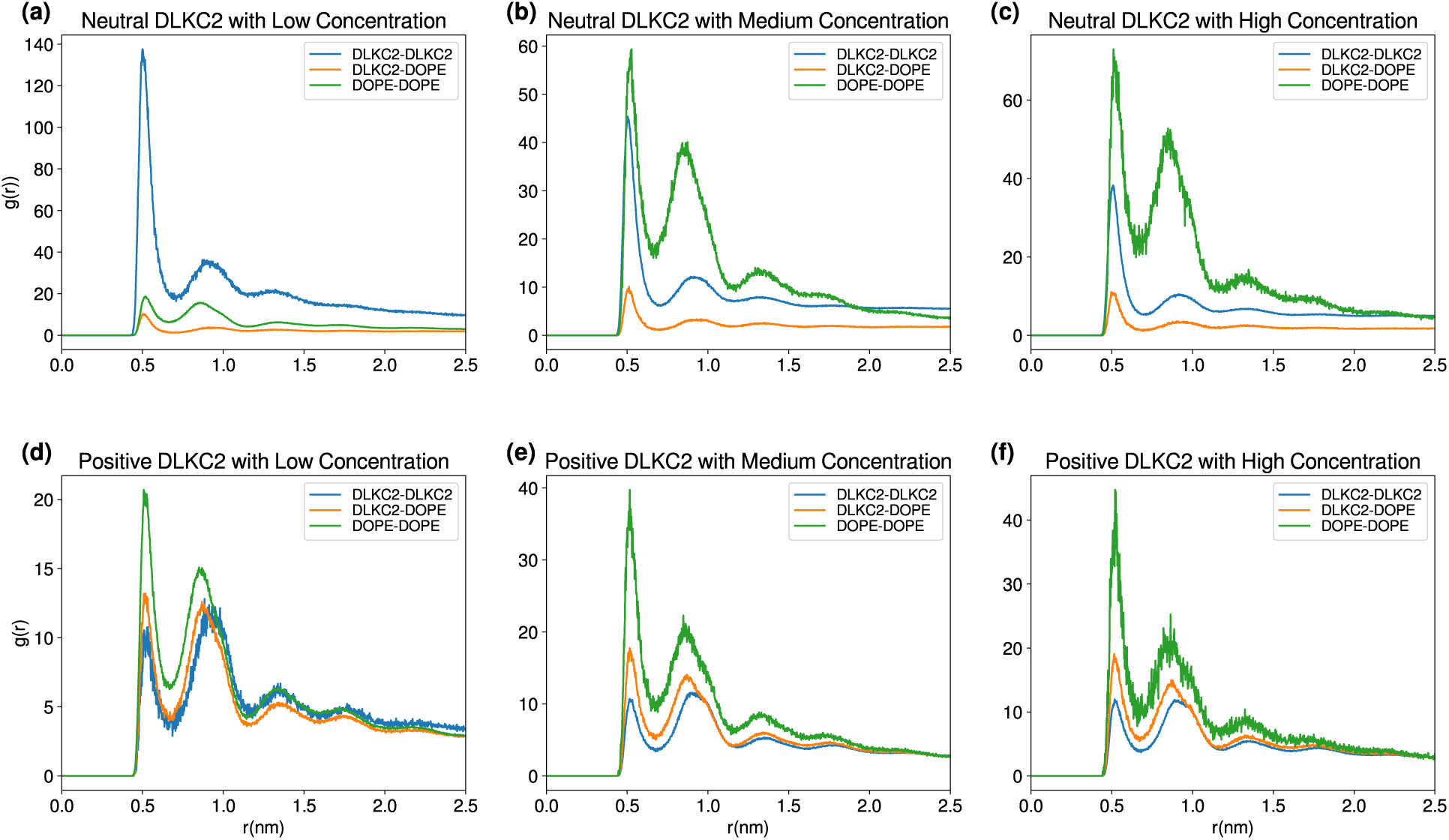
Radial distribution function of lipids to understand the distribution of individual lipids in the LNPs.

**Figure 7:**
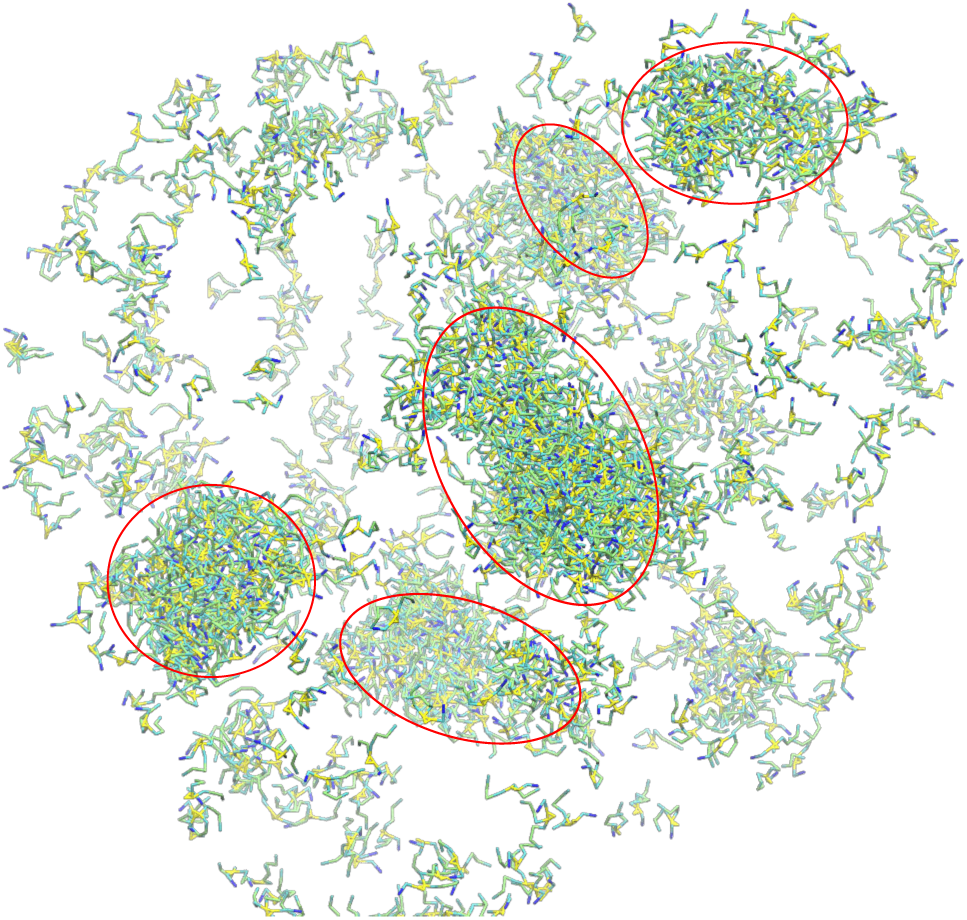
Clustering of DLKC2-DLCK2 in the LNP with low concentration of neutral DLKC2 is shown.

**Figure 8:**
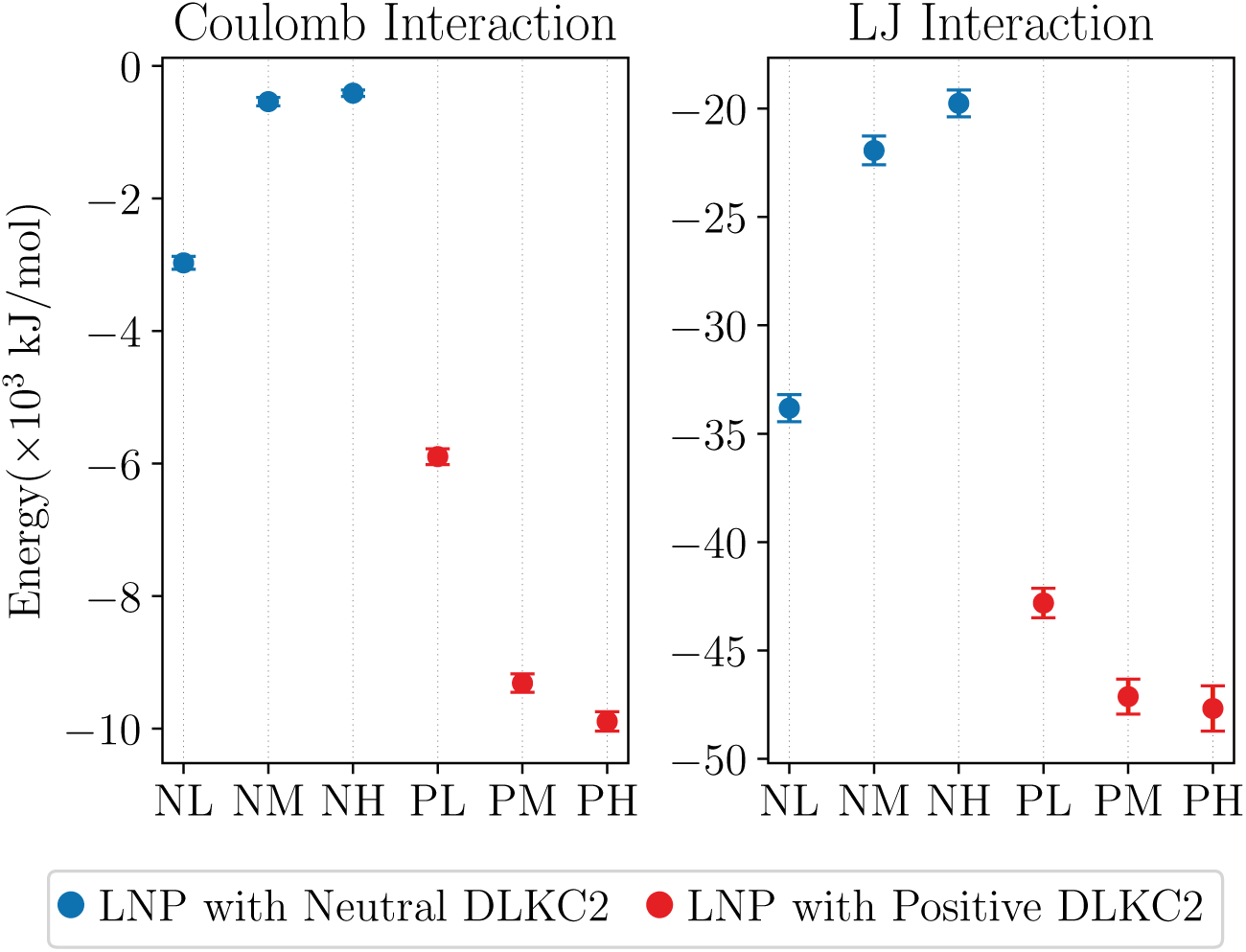
Short range LJ and Coulomb potential interaction energy between RNA and lipids. NL, NM, and NH, represent neutral DLKC2 with low, medium and high concentrations in LNP. PL, PM, and PH, represent cationic DLKC2 with low, medium and high concentrations in LNP.

### Force experience by siRNA while crossing membrane

In addition to the self assembly of LNP for the encapsulation of siRNA, understanding the escape of siRNA from lipid nanoparticles (LNPs) is crucial to comprehend the stability of the formulation. As mentioned in the methods section, an all-atom bilayer system was prepared to study the steered molecular dynamics simulation. However, before conducting a SMD simulation on an all-atom bilayer system, the simulation was initially performed on the LNP with a medium concentration of positive DLKC2. In this system, one of the siRNA close to the membrane was tried pulling from inside to outside of the LNP, as shown in the Figure 9. Even after 20 ns of pulling, the RNA never came out of the membrane; instead, it started pushing the membrane in the direction of pulling. This is due to the less degrees of freedom and bulkier size of the coarse-grained model. Therefore, to overcome this problem and gain a detailed insights into the interactions between lipids and siRNA, an all-atom bilayer was considered as a a more detailed model.

**Figure 9:**
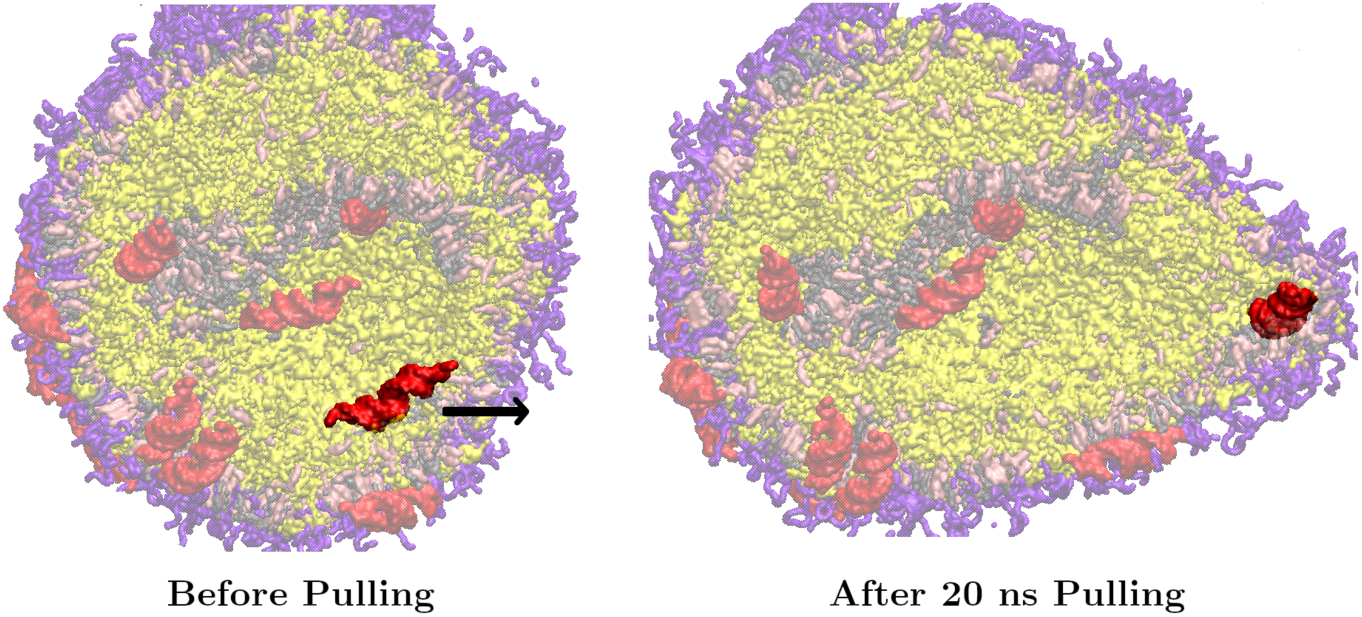
Pulling of the siRNA (highlighted red color) was conducted for the steered MD simulation in the direction specified by arrow in the CG model. After 20 ns of pulling, the RNA made a bulge instead of passing through the membrane.

The SMD simulation was performed on the siRNA by pulling the center of mass of siRNA along the negative z-direction to cross the bilayer. During the pulling of the siRNA through the lipid bilayer, it was observed that due to the presence of positive ionizable lipids, the negatively charged siRNA experiences a strong force to pass through the positively charged lipid layer (Figure 10). It can be seen from Figure 11a that the membrane gets deformed while the siRNA is pulled out to the lower side. The head group of the lipids experiences a strong attraction towards the siRNA, which can be seen from Figure 12. Such effect is not observed in the membrane with neutral DLKC2 since the siRNA could cross the bilayer without affecting the membrane layer, as demonstrated in Figure 11b. Using this method, it can be verified that while encapsulating siRNA, the positive ionizable lipid plays a significant role and helps in protecting the siRNA inside the vesicles. Figure 10 shows the restoring force experienced by siRNA while pulling in positive and neutral membrane cases. The distance is measured between the center of mass of siRNA and center of mass of lipid bilayer. Negative value of distance represents negative z direction with respect to the center of mass of lipid bilayer. For the neutral lipid, the force restores back to the starting value, whereas in the case of the positive lipid, the force doesn’t go to zero.

**Figure 10:**
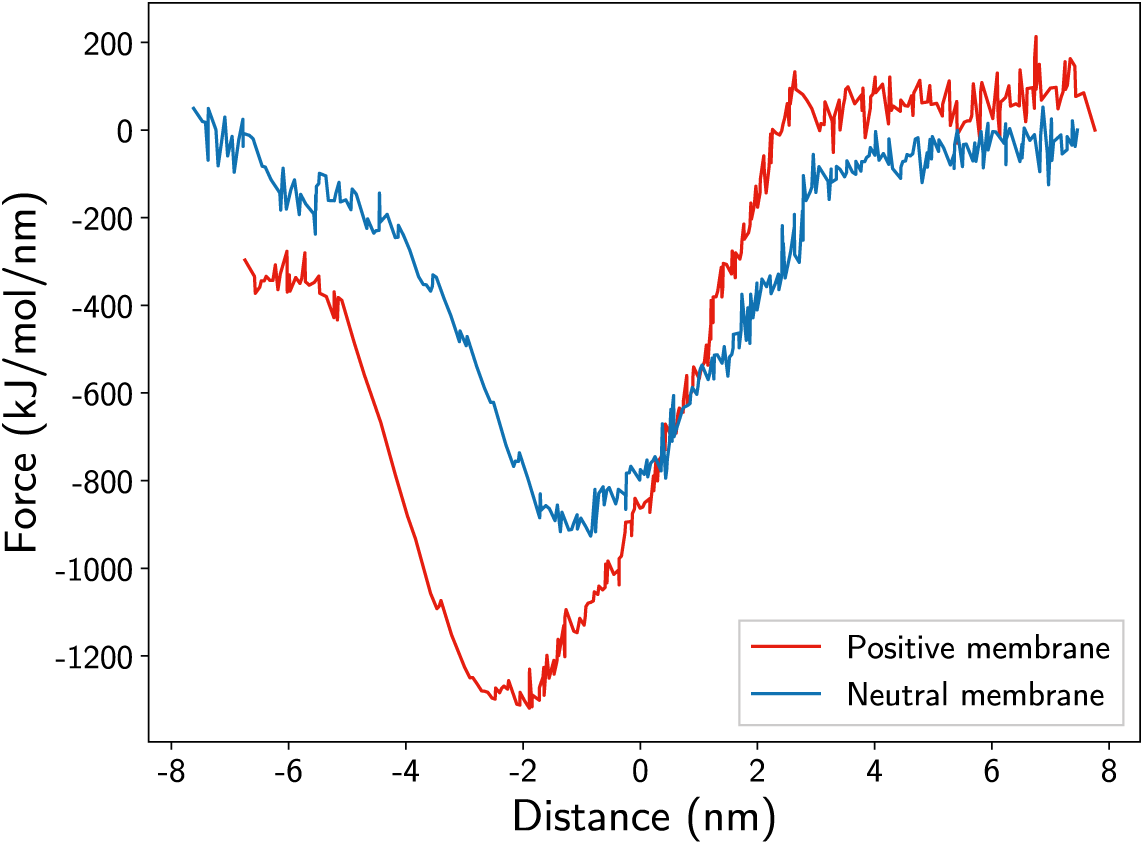
Restoring force experienced by the siRNA while crossing the membrane. Distance is measured between center of mass of siRNA and the center of mass of lipid bilayer.

**Figure 11:**
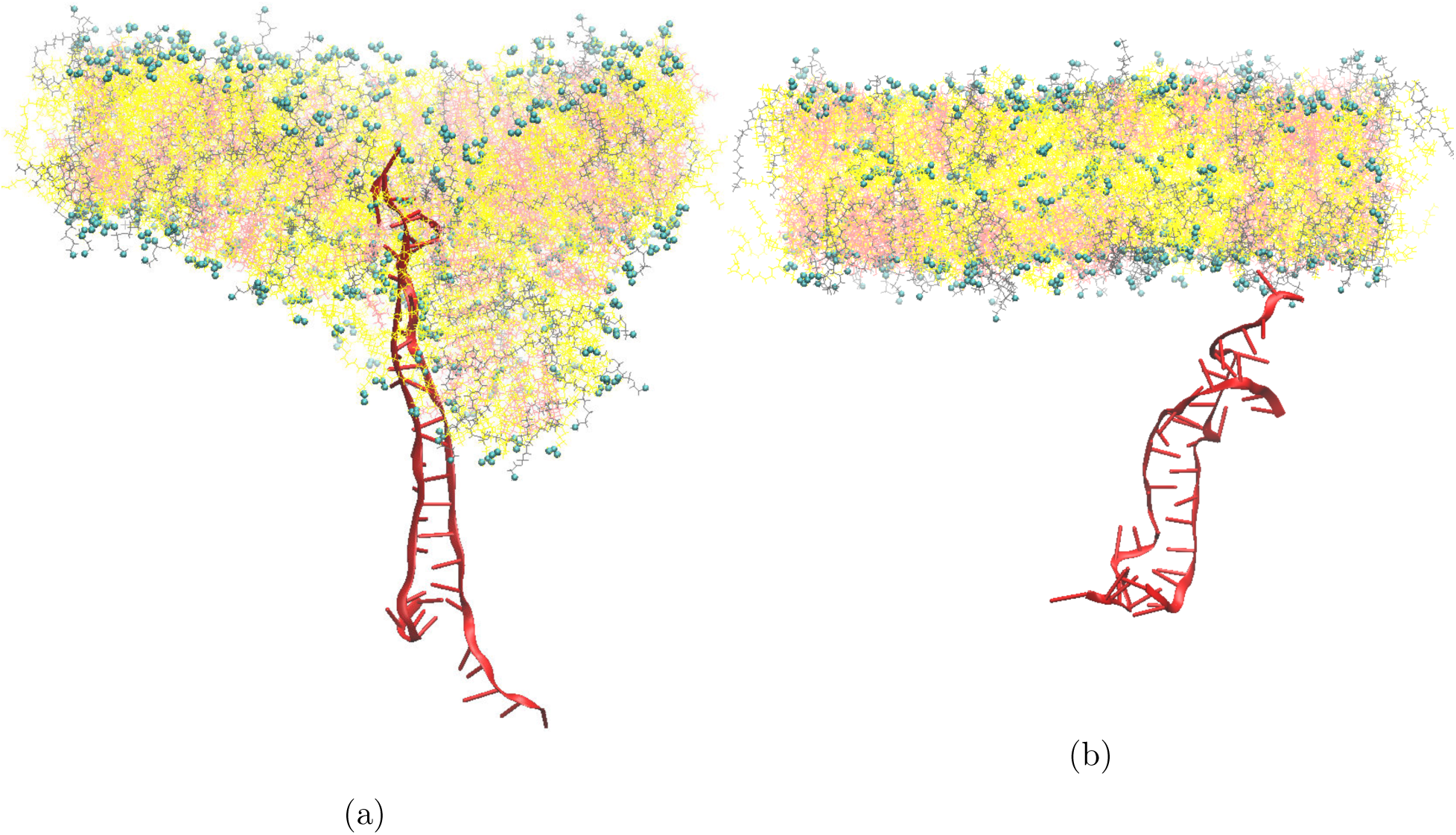
During the end of pulling simulation in bilayers containing (a) positive DLKC2, and (b) neutral DLKC2. Due to strong attraction between negative siRNA and positive DLKC2, the membrane gets deformed while the siRNA start passing through. DLKC2 is shown in yellow, cholesterol in pink, DOPE in gray, lipid head group in cyan, and RNA in red; water is not shown for clarity.

**Figure 12:**
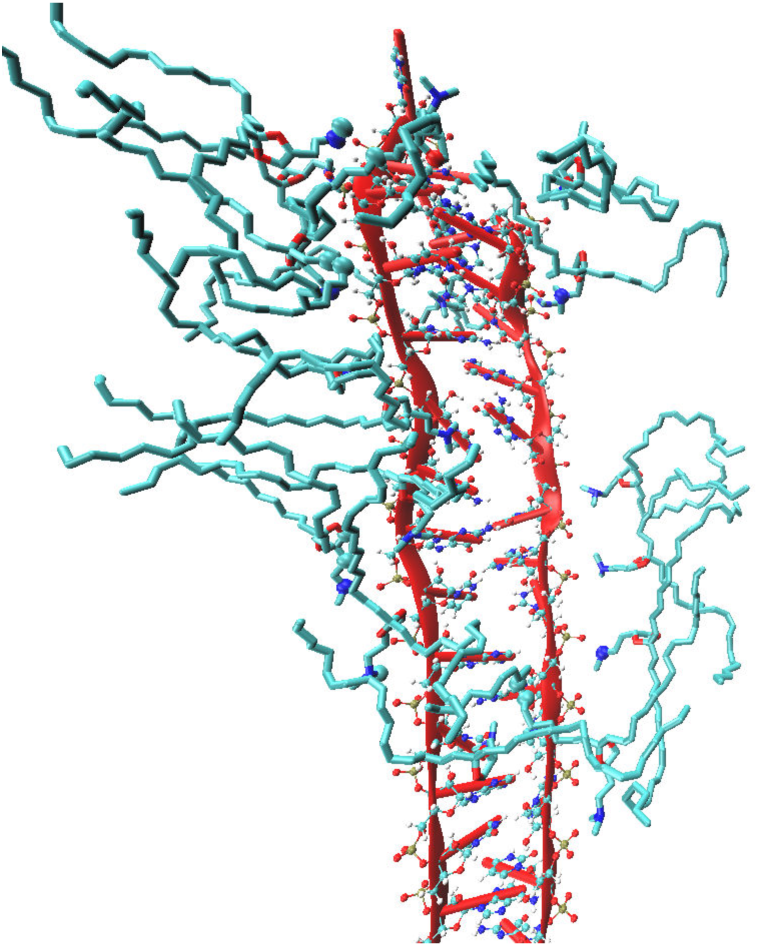
Zoomed image from Figure 11a shows that head group of DLKC2 lipids which are positive, are attached to the negative phosphate group of siRNA strands.

## Conclusion

In the current study, we demonstrated the effects of various concentrations of ionizable lipids (DLKC2) and phospholipids (DOPE) along with cholesterol and PEGlyted lipids on the stability of the LNPs and encapsulation of RNAs. We found that the LNPs containing positive DLKC2 with low and medium concentrations are able to encapsulate all the siRNA, which is consistent with the previous experimental and simulation studies.^41^ The study also found that neutral DLKC2 with low concentrations showed similar LNP formation to positive DLKC2 encapsulating all the siRNA, but not with medium and higher concentrations of neutral DLKC2. In these two cases, only 25% of the siRNAs were encapsulated. We observed the formation of many compartments inside the LNPs where the siRNAs were found to affect the encapsulation of siRNA. The higher the number of compartments, the higher the number of siRNA encapsulated. In the case of medium and high concentrations of neutral DLKC2, only two compartments were seen, whereas in positive DLKC2, at least five were seen. As the concentration of neutral DLKC2 increases in the LNPs, a distinct lipid bilayer is not observed. As a result, we do not observe many compartments inside the LNPs. A similar effect is seen for LNPs with very high concentrations of positive DLKC2. As for the previous studies,^33^ the presence of positive lipids is important for the stabilization and resistance to degradation of nucleic acids. Due to the negative charge of the siRNAs, a stronger interaction towards the lipid head group was observed, which facilitated the encapsulation. For the first time, this study compared the mechanism of encapsulation of RNAs inside positively charged and neutral lipid nanoparticles with varying lipid concentration using a molecular dynamics study. Although the LNP with positively charged lipids is crucial for encapsulating RNAs, we also found that the composition of lipids with a low concentration of DLKC2 and a high concentration of DOPE can be good for forming LNPs. But, in this system, many areas with local aggregates of DLKC2 are found in the lipid bilayer in the LNP. This may lead to the instability of LNP in the later stage of drug delivery. The force experience by siRNA calculated using steered MD simulation to cross the bilayer containing positive ionizable lipids was 400 kJ/mol/nm more than that of bilayer with neutral ionizable lipids. This suggests that the formation of encapsulated siRNA is more favourable with positive ionizable lipids. Our work was limited to the stability of siRNAs inside LNP with various lipid compositions. While we acknowledge the significance of endosomal escape in the area of RNA-based therapeutics, we do not discuss the endosomal escape as it falls outside the scope of our investigation. Our primary focus was to elucidate the impact of positive and neutral ionizable lipids on the stability of LNPs during the formulation process. Specifically, the process of siRNA encapsulation in the presence of positive and neutral ionizable lipids. However, our study provides valuable insights into the development of RNA therapeutics and the use of LNPs as a delivery system. Further studies are needed to complete the understanding of the delivery and endosomal escape of RNA-based drugs.

## Author Contributions

MRB: conceptualization, methodology, analysis, writing - review & editing. SR: conceptualization, investigation, methodology, writing - review & editing. JKS: investigation, writing - review & editing.

## Acknowledgements

The authors thank the support and the resources provided by PARAM Sanganak under the National Supercomputing Mission, Government of India at the Indian Institute of Technology, Kanpur. MRB thank Indian Institution of Technology Kanpur for the funding.

## Supporting Information

Following information is provided in the supporting information.

- CG Martini 2 bead types of siRNA, DOPE, Cholesterol, PEG-lipids
- Initial configuration of steered MD simulation
- Distribution of DLKC2 in the neutral and postive LNP with low concentration of DLKC2
- siRNA - lipid interaction energy

