## Supporting Information for "*In Silico* Engineering of Stable siRNA Lipid Nanoparticles: Exploring the Impact of Ionizable Lipid Concentrations for Enhanced Formulation Stability"

### 1 siRNA

All atom structure of siRNA was take from PDB id, 2F8S. The sequence of the siRNA is given by: `AGACAGCAUAUAUGCUGUCUUU`

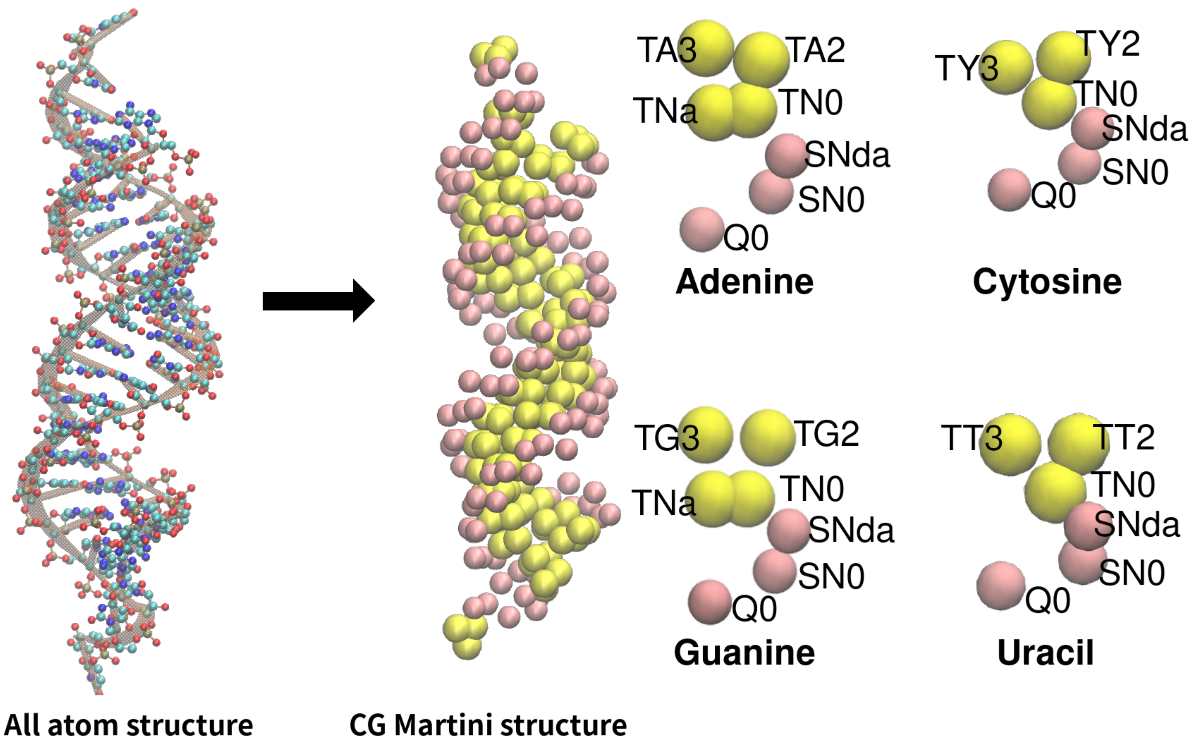


Figure S1: CG Martini structure of siRNA. The bead types are according to martini version 2 convention. Yellow beads are related to backbone and pink beads are related to side chain. Martini bead types of adenine, guanine, cytosine, and uracil are also provided.

### 2 PEG

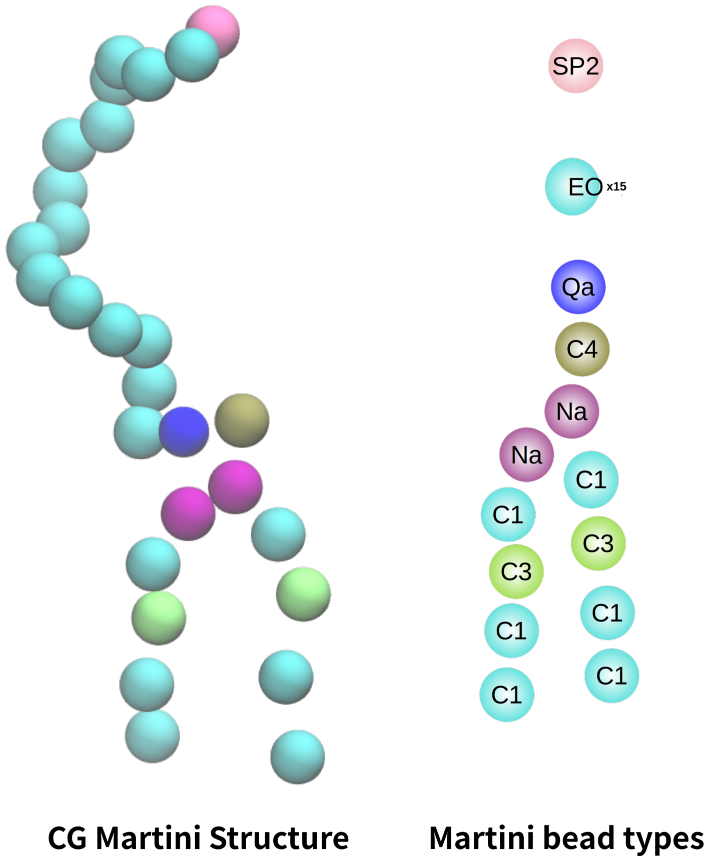


Figure S2: CG Martini structure of PEG. The bead types are according to martini version 2 convention.

### 3 DOPE

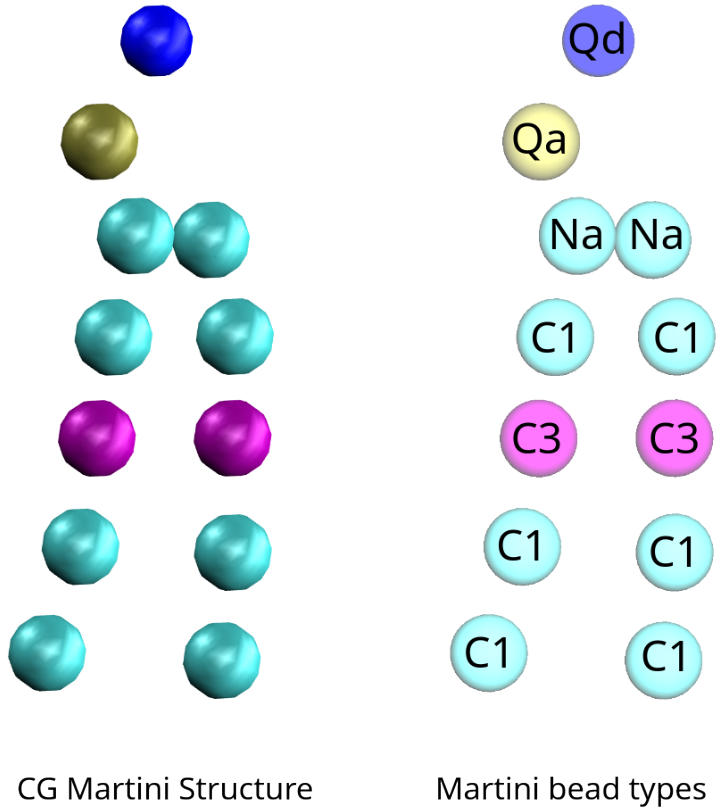


Figure S3: CG Martini structure of DOPE. The bead types are according to martini version 2 convention.

### 4 Cholesterol

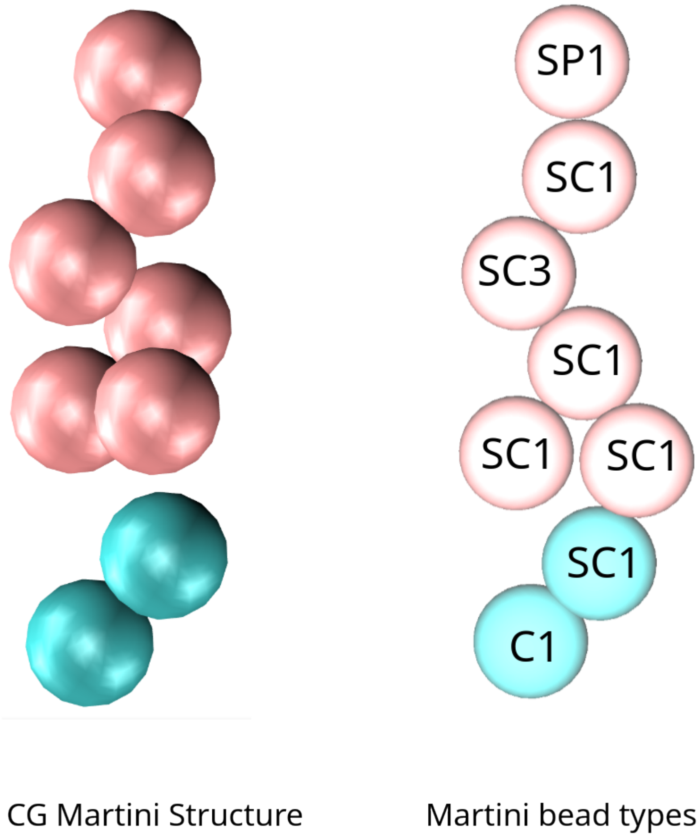


Figure S4: CG Martini structure of cholesterol. The bead types are according to martini version 2 convention.

### 5 Steered molecular dynamics study

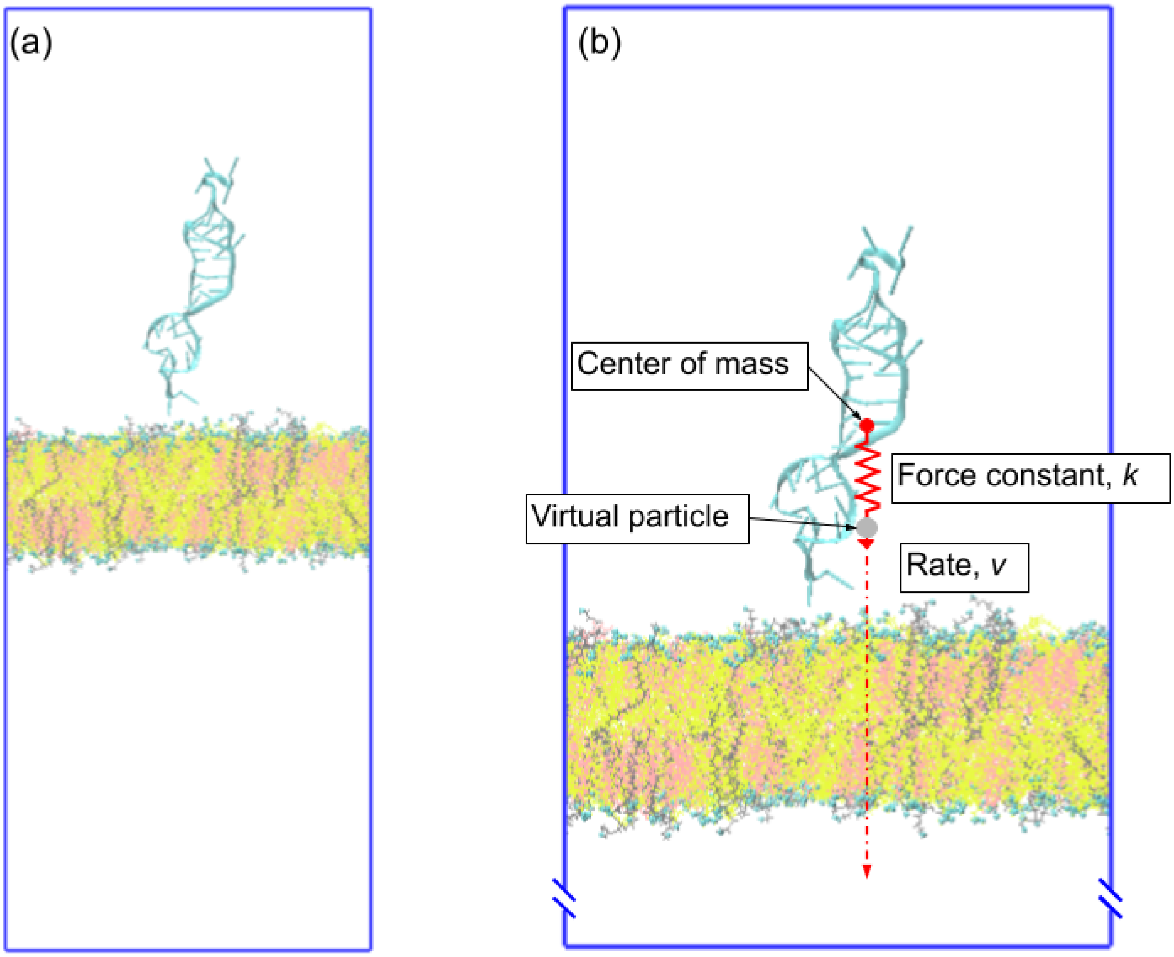


Figure S5: (a)Starting structure of steered molecular dynamics simulation. (b)An external harmonic potential, $U' = 1/2k(x_0 - x - vt)^2$, is applied to the centre of mass of siRNA with a force constant of $k$ =600$kJ/mol/nm^2$ applied between the center of mass of siRNA and a virtual particle moving along the negative z-direction with a constant velocity of $v$ = 0.5 ns per nm. $x_0$ is the initial position of the restraint point.

### 6 DLKC2 distribution

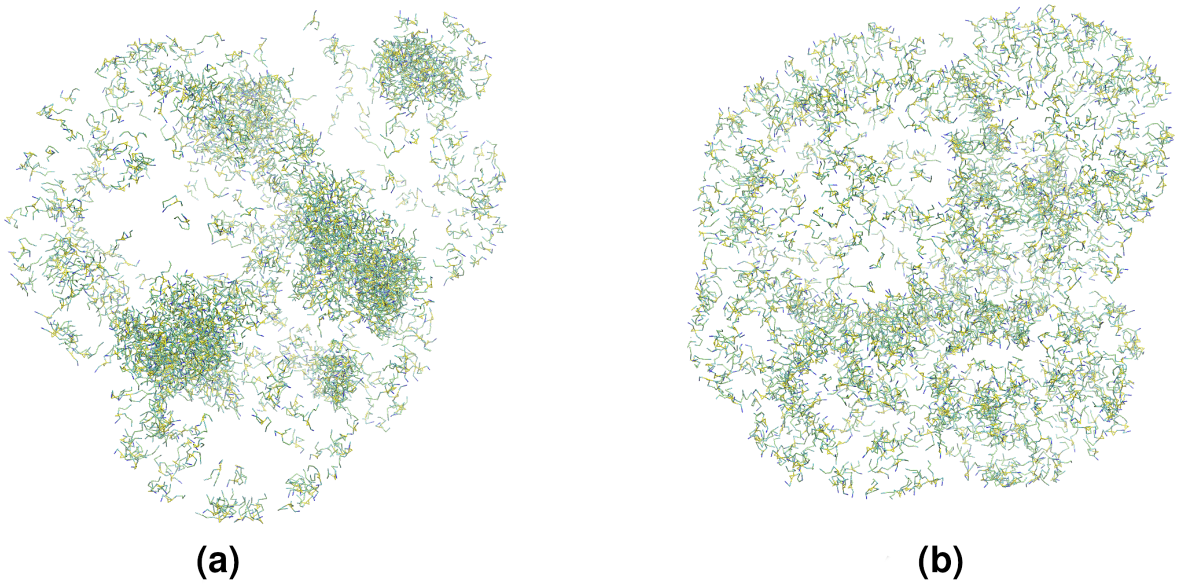


Figure S6: Distribution of DLKC2 in LNP with (a) low concentration of neutral DLKC2 and (b) low concentration of positive DLKC2. (a) In LNP with neutral ionisable lipids, the DLKC2 molecules are seen aggregated, (b) but in case of positive DLKC2, the lipids are equally distributed.

### 7 siRNA - lipid interaction energy

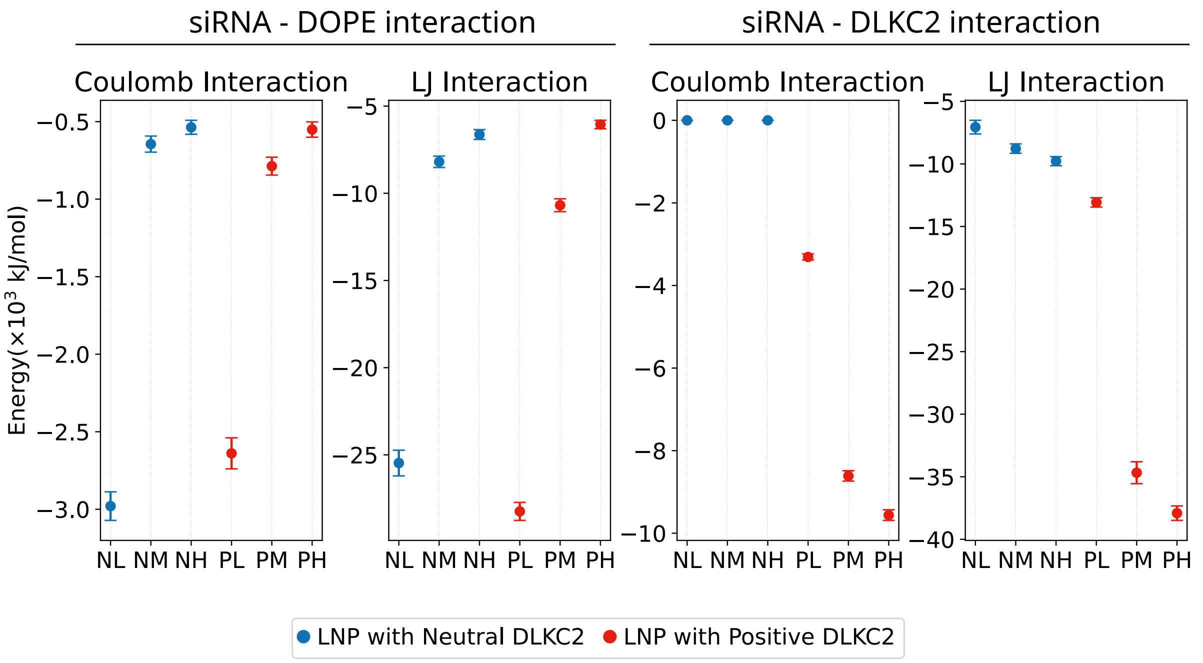


Figure S7: Short range LJ and coulomb potential interaction energy between siRNA - DOPE, and siRNA - DLKC2 . NL, NM, and NH, represent neutral DLKC2 with low, medium and high concentrations in LNP. PL, PM, and PH, represent cationic DLKC2 with low, medium and high concentrations in LNP.

### 8 Compartments in LNPs where siRNAs are encapsulated

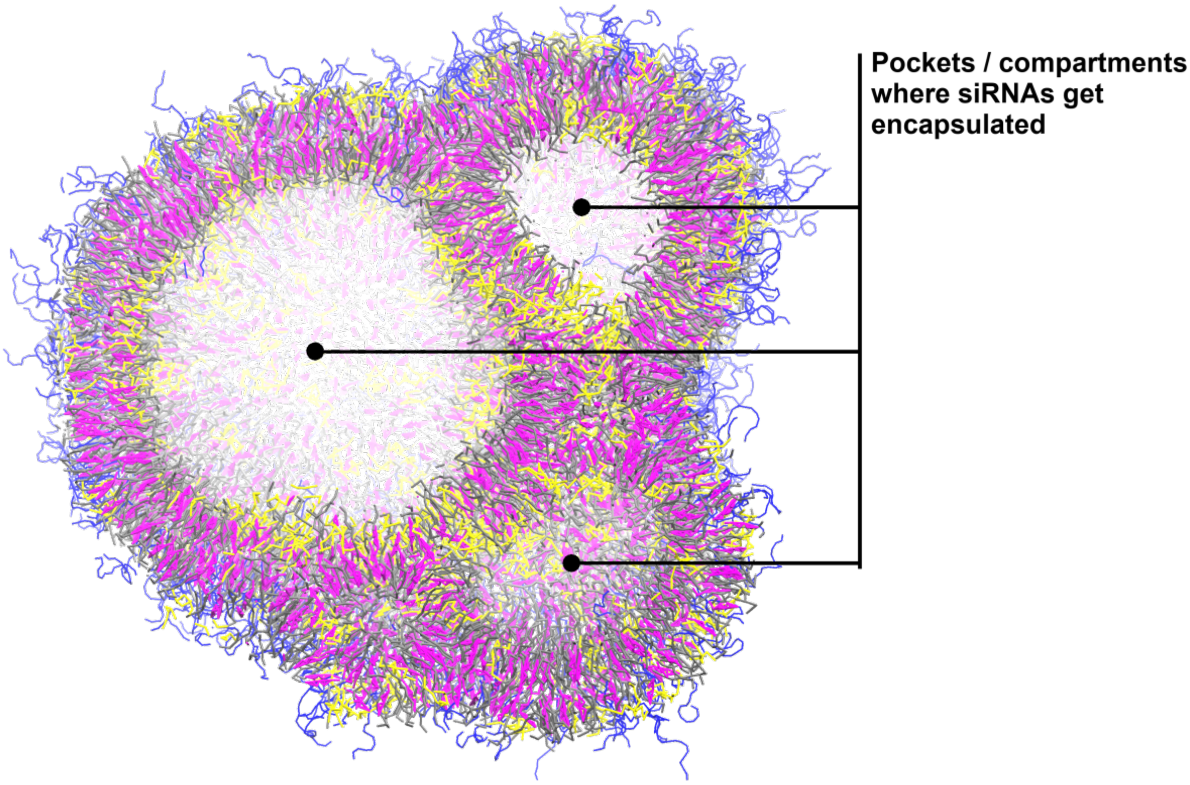


Figure S8: Schematic of LNP showing pockets or compartments where siRNAs are encapsulated.
